# Mapping Cortical Brain Asymmetry in 17,141 Healthy Individuals Worldwide via the ENIGMA Consortium

**DOI:** 10.1101/196634

**Authors:** Xiang-Zhen Kong, Samuel R. Mathias, Tulio Guadalupe, Christoph Abé, Ingrid Agartz, Theophilus N. Akudjedu, Aleman Andre, Alhusaini Saud, Nicholas B. Allen, David Ames, Ole A. Andreassen, Alejandro Arias Vasquez, Nicola J. Armstrong, Felipe Bergo, Mark E. Bastin, Albert Batalla, Jochen Bauer, Bernhard T Baune, Ramona Baur, Joseph Biederman, Sara K. Blaine, Premika Boedhoe, Erlend Bøen, Anushree Bose, Janita Bralten, Daniel Brandeis, Silvia Brem, Henry Brodaty, Henrieke Bröhl, Samantha J. Brooks, Jan Buitelaar, Christian Bürger, Robin Bülow, Vince Calhoun, Anna Calvo, Erick Jorge Canales-Rodríguez, Jose M. Canive, Dara M. Cannon, Elisabeth C. Caparelli, Francisco X. Castellanos, Gianpiero L. Cavalleri, Fernando Cendes, Tiffany Moukbel Chaim-Avancini, Kaylita Chantiluke, Qun-lin Chen, Xiayu Chen, Yuqi Cheng, Anastasia Christakou, Vincent P. Clark, David Coghill, Colm G. Connolly, Annette Conzelmann, Aldo Cόrdova-Palomera, Janna Cousijn, Tim Crow, Ana Cubillo, Udo Dannlowski, Sara Ambrosino de Bruttopilo, Patrick de Zeeuw, Ian J. Deary, Norman Delanty, Damion V. Demeter, Adriana Di Martino, Erin W Dickie, Bruno Dietsche, N. Trung Doan, Colin P. Doherty, Alysa Doyle, Sarah Durston, Eric Earl, Stefan Ehrlich, Carl Johan Ekman, Torbjørn Elvsåshagen, Jeffery N. Epstein, Damien A. Fair, Stephen Faraone, Helena Fatouros-Bergman, Guillen Fernndez, Geraldo Busatto Filho, Lena Flyckt, Katharina Forster, Fouche Jean-Paul, John J. Foxe, Paola Fuentes-Claramonte, Janice Fullerton, Hugh Garavan, Danielle do Santos Garcia, Ian H. Gotlib, Anna E. Goudriaan, Hans Jorgen Grabe, Nynke A. Groenewold, Dominik Grotegerd, Oliver Gruber, Tiril Gurholt, Jan Haavik, Tim Hahn, Narelle K. Hansell, Mathew A. Harris, Catharina Hartman, Maria del Carmen Valdes Hernandez, Dirk Heslenfeld, Robert Hester, Derrek Paul Hibar, Beng-Choon Ho, Tiffany C. Ho, Pieter J. Hoekstra, Ruth J. van Holst, Martine Hoogman, Marie F. Hovik, Fleur M. Howells, Kenneth Hugdahl, Chaim Huyser, Martind Ingvar, Lourdes Irwin, Akari Ishikawa, Anthony James, Neda Jahanshad, Terry L. Jernigan, Erik G Jonsson, Claas Kahler, Vasily Kaleda, Clare Kelly, Michael Kerich, Matcheri S Keshavan, Sabin Khadka, Tilo Kircher, Gregor Kohls, Kerstin Konrad, Ozlem Korucuoglu, Bernd Kramer, Axel Krug, Jun Soo Kwon, Nanda Lambregts-Rommelse, Mikael Landen, Luisa Lazaro, Irina Lebedeva, Rhoshel Lenroot, Klaus-Peter Lesch, Qinqin Li, Kelvin O. Lim, Jia Liu, Christine Lochner, Edythe D. London, Vera Lonning, Valentina Lorenzetti, Michelle Luciano, Maartje Luijten, Astri J. Lundervold, Scott Mackey, Frank P. MacMaster, Sophie Maingault, Charles B. Malpas, Ulrik F. Malt, David Mataix-Cols, Rocio Martin-Santos, Andrew R. Mayer, Hazel McCarthy, Philip B. Mitchell, Bryon A. Mueller, Susana Munoz Maniega, Bernard Mazoyer, Colm McDonald, Quinn McLellan, Katie L. McMahon, Genevieve McPhilemy, Reza Momenan, Angelica M. Morales, Janardhanan C. Narayanaswamy, Jose Carlos Vasques Moreira, Stener Nerland, Liam Nestor, Joel T. Nigg, Jan-Egil Nordvik, Stephanie Novotny, Eileen Oberwelland, Ruth L. O'Gorman, Jaap Oosterlaan, Bob Oranje, Catherine Orr, Bronwyn Overs, Paul Pauli, Martin Paulus, Kerstin Plessen, Georg G. von Polier, Edith Pomarol-Clotet, Jiang Qiu, Joaquim Radua, Josep Antoni Ramos-Quiroga, Y.C. Janardhan Reddy, Andreas Reif, Gloria Roberts, Pedro Rosa, Katya Rubia, Matthew D. Sacchet, Perminder S. Sachdev, Raymond Salvador, Lianne Schmaal, Lisanne Schweren, Larry Seidman, Jochen Seitz, Mauricio Henriques Serpa, Philip Shaw, Elena Shumskaya, Timothy J. Silk, Alan N. Simmons, Egle Simulionyte, Rajita Sinha, Zsuzsika Sjoerds, Runar Elle Smelror, Joan Carlos Soliva, Nadia Solowij, Scott R. Sponheim, Dan J. Stein, Elliot A. Stein, Michael Stevens, Lachlan T. Strike, Gustavo Sudre, Jing Sui, Leanne Tamm, Hendrik S. Temmingh, Robert J. Thoma, Alexander Tomyshev, Giulia Tronchin, Jessica Turner, Anne Uhlmann, Theo G.M. van Erp, Odile van den Heuvel, Dennis van der Meer, Liza van Eijk, Alasdair Vance, Ilya M. Veer, Dick J. Veltman, Ganesan Venkatasubramanian, Oscar Vilarroya, Yolanda Vives-Gilabert, Aristotle N. Voineskos, Henry Volzke, Daniella Vuletic, Susanne Walitza, Henrik Walter, Esther Walton, Joanna M. Wardlaw, Wei Wen, Lars T. Westlye, Christopher D. Whelan, Tonya White, Reinout W. Wiers, Margaret J. Wright, Katharina Wittfeld, Tony T. Yang, Clarissa L. Yasuda, Yuliya Yoncheva, Murat Yucel, Je-Yeon Yun, Marcus Vinicius Zanetti, Zonglei Zhen, Xing-xing Zhu, Georg C. Ziegler, Kathrin Zierhut, Greig I. de Zubicaray, Marcel Zwiers, Karolinska Schizophrenia Project KaSP, David C. Glahn, Barbara Franke, Fabrice Crivello, Nathalie Tzourio-Mazoyer, Simon E. Fisher, Paul M. Thompson, Clyde Francks, Lars Farde, Goran Engberg, Sophie Erhardt, Simon Cervenka, Lilly Schwieler, Fredrik Piehl, Karin Collste, Pauliina Victorsson, Anna Malmqvist, Mikael Hedberg, Funda Orhan

**Affiliations:** Language and Genetics Department, Max Planck Institute for Psycholinguistics, Nijmegen, The Netherlands.; Department of Psychiatry, Yale School of Medicine, New Haven, CT 06519, USA.; Department of Clinical Neuroscience, Osher Centre, Karolinska Institutet, Stockholm, Sweden.; Norwegian Centre for Mental Disorders Research (NORMENT), K. G. Jebsen Centre for Psychosis Research, Institute of Clinical Medicine, University of Oslo, Oslo, Norway.; Department of Clinical Neuroscience, Centre for Psychiatry Research, Karolinska Institutet, Stockholm, Sweden.; The Centre for Neuroimaging & Cognitive Genomics (NICOG), Clinical Neuroimaging Lab, NCBES Galway Neuroscience Centre, College of Medicine, Nursing, and Health Sciences, National University of Ireland Galway, H91 TK33 Galway Ireland, Republic of Ireland.; BCN Neuroimaging Center, Department of Neuroscience, University Medical Center Groningen, University of Groningen, The Netherlands.; Department of Molecular and Cellular Therapeutics, Royal College of Surgeons in Ireland, Dublin, Ireland.; Neurology and Neurosurgery Department, Montreal Neurological Hospital and Institute, McGill University, Montreal, Canada.; Orygen, The National Centre of Excellence in Youth Mental Health, Parkville, Australia.; Department of Psychology, University of Oregon, Eugene OR, USA.; National Ageing Research Institute, Melbourne, Australia.; Academic Unit for Psychiatry of Old Age, University of Melbourne, Melbourne, Australia.; Norwegian Centre for Mental Disorders Research (NORMENT), KG Jebsen Centre for Psychosis Research, Division of Mental Health and Addiction, Oslo University Hospital & Institute of Clinical Medicine, University of Oslo, Oslo, Norway.; Department of Human Genetics, Radboud University Medical Center, Nijmegen, The Netherlands.; Department of Psychiatry, Radboud University Medical Center, Nijmegen, The Netherlands.; Department of Cognitive Neuroscience, Radboud University Medical Center, Nijmegen, The Netherlands.; Donders Institute for Brain, Cognition and Behaviour, Radboud University, Nijmegen, The Netherlands.; Mathematics and Statistics, Murdoch University, Perth, Australia.; Laboratory of Neuroimaging, Department of Neurology, University of Campinas.; Centre for Clinical Brain Sciences, University of Edinburgh, Edinburgh, UK.; Brain Research Imaging Centre, University of Edinburgh, Edinburgh, UK.; Department of Psychiatry, Donders Institute for Brain, Cognition and Behaviour, Radboud University Medical Centre, Nijmegen, The Netherlands.; Department of Clinical Radiology, School of Medicine, University of Münster, Germany.; Discipline of Psychiatry, University of Adelaide, Australia.; Department of Psychology, University of Würzburg, Germany, Würzburg, Germany.; Department of Psychiatry, Harvard Medical School, Boston, Mass, USA.; Clinical and Research Programs in Pediatric Psychopharmacology and Adult ADHD, Massachusetts General Hospital, Boston, MA, USA.; Department of Psychiatry, Yale University School of Medicine, New Haven, CT, USA.; Department of Psychiatry, VU University Medical Center, Amsterdam, The Netherlands.; Department of Anatomy & Neurosciences, VU University Medical Center, Amsterdam, The Netherlands.; Amsterdam Neuroscience, Amsterdam, The Netherlands.; Department of Psychiatric Research, Diakonhjemmet Hospital, Oslo, Norway.; Department of Psychiatry, National Institute of Mental Health and Neurosciences, Bengaluru, India.; Donders Institute for Brain, Cognition and Behaviour, Nijmegen, The Netherlands.; University Clinics for Child and Adolescent Psychiatry (UCCAP), University of Zurich, Zurich, Switzerland.; Neuroscience Center Zurich, University of Zurich and ETH Zurich, Switzerland.; Zurich Center for Integrative Human Physiology, University of Zurich, Zurich, Switzerland.; Department of Child and Adolescent Psychiatry and Psychotherapy, Central Institute of Mental Health, Medical Faculty Mannheim/Heidelberg University, J5, 68159 Mannheim, Germany.; Centre for Healthy Brain Ageing, School of Psychiatry, UNSW Australia.; Dementia Collaborative Research Centre ØC Assessment and Better Care, University of New South Wales, Sydney, Australia.; Department of Psychiatry and Psychotherapy Philipps-University Marburg, Germany.; Department of Psychiatry and Mental Health, University of Cape Town, South Africa.; Karakter Child and Adolescent Psychiatry, Nijmegen, The Netherlands.; Department of Cognitive Neuroscience, Radboud university, Nijmegen, The Netherlands.; Department of Psychiatry, University of Münster, Germany.; Department of Diagnostic Radiology and Neuroradiology, University Medicine Greifswald, Greifswald, Germany.; The Mind Research Network, Albuquerque, NM, USA.; Department of Electrical and Computer Engineering, University of New Mexico, Albuquerque, NM 87131, United States.; Medical Image Core Facility, August Pi I Sunyer Biomedical Research Institute (IDIBAPS), Barcelona, Spain.; CIBERBBN.; FIDMAG Germanes Hospitalaries Research Foundation, Barcelona, Spain.; CIBERSAM, Centro de Investigatión Biomédica en Red de Salud Mental, Spain.; Departments of Psychiatry and Neurosciences, University of New Mexico.; Neuroimaging Research Branch, National Institute on Drug Abuse, National Institutes of Health, Baltimore, Maryland, USA.; The Child Study Center at Hassenfeld Children's Hospital at NYU Langone, New York, USA.; Division of Child and Adolescent Psychiatric Research, Nathan Kline Institute for Psychiatric Research, Orangeburg, NY, USA.; Department of Psychiatry, Faculty of Medicine, University of São Paulo, São Paulo, Brazil.; Center for Interdisciplinary Research on Applied Neurosciences (NAPNA), University of São Paulo, São Paulo, Brazil.; King's College London, Institute of Psychiatry, Psychology and Neuroscience, Department of Child and Adolescent Psychiatry, London, UK.; School of Psychology, Southwest University, Chongqing, China.; Key Laboratory of Cognition and Personality, Ministry of Education, Chongqing, China.; State Key Laboratory of Cognitive Neuroscience and Learning & IDG/McGovern Institute for Brain Research, Faculty of Psychology, Beijing Normal University, Beijing, China.; Department of Psychiatry, First Affiliated Hospital of Kunming Medical University, Kunming, China.; Department of Child and Adolescent Psychiatry, Institute of Psychiatry, King's College London, London WC2R 2LS, UK.; School of Psychology and Clinical Language Sciences, University of Reading, Reading RG6 6AL, UK.; Department of Psychology, University of New Mexico, Albuquerque, NM 87131, United States.; Departments of Paediatrics and Psychiatry, University of Melbourne. Victoria, Australia.; Division of Neuroscience, Ninewells Hospital and Medical School, University of Dundee.; Department of Biomedical Sciences, Florida State University, Tallahassee, FL 32306, USA.; Department of Psychiatry, Division of Child and Adolescent Psychiatry, and Weill Institute for Neurosciences, University of California, San Francisco, 401 Parnassus Avenue, San Francisco, CA, USA.; Department of Child and Adolescent Psychiatry and Psychotherapy, University of Tübingen, Tübingen,Germany.; Norwegian Centre for Mental Disorder Research (NORMENT), K.G. Jebsen Centre for Psychosis Research, Division of Mental Health and Addiction, Oslo University Hospital & Institute of Clinical Medicine, University of Oslo, Oslo, Norway.; Department of Developmental Psychology, University of Amsterdam, Amsterdam, The Netherlands.; SANE POWIC, University Department of Psychiatry, Warneford Hospital, Oxford, UK.; NICHE-lab, Brain Center Rudolf Magnus, Department of Psychiatry, University Medical Center Utrecht, Utrecht, The Netherlands.; Centre for Cognitive Ageing and Cognitive Epidemiology, Psychology, University of Edinburgh, Edinburgh, UK.; Department of Behavioral Neuroscience at Oregon Health & Science University, Portland, OR, USA.; Kimel Family Translational Imaging-Genetics Research Laboratory, Campbell Family Mental Health Research Institute, Center for Addiction and Mental Health, Toronto, Canada.; Neurology Department, St. James's Hospital, Dublin, Ireland.; Department of Psychiatry & Center for Genomic Medicine, Massachusetts General Hospital, Harvard Medical School, Boston, MA,USA.; Stanley Center for Psychiatric Research at the Broad Institute, Cambridge, MA, USA.; Division of Psychological and Social Medicine and Developmental Neurosciences, Faculty of Medicine, Technische Universitat Dresden, Dresden, Germany.; Norwegian Centre for Mental Disorder Research (NORMENT), Institute of Clinical Medicine, University of Oslo, Oslo, Norway.; Department of Neurology, Oslo University Hospital, Oslo, Norway.; University of Cincinnati College of Medicine, Cincinnati, OH, USA.; Cincinnati Children's Hospital Medical Center, Cincinnati, OH, USA.; Department of Behavioral Neuroscience, Oregon Health & Science University, USA.; Department of Psychiatry, Oregon Health & Science University, USA.; Advanced Imaging Research Center, Oregon Health & Science University, USA.; K.G. Jebsen Centre for Neuropsychiatric Disorders, Department of Biomedicine, University of Bergen, Bergen, Norway.; Department of Psychiatry, SUNY Upstate Medical University, Syracuse, NY, USA.; Donders Institute for Brain, Cognition and Behaviour, Radboud University Medical Center, Nijmegen, The Netherlands.; Institute of Psychiatric Phenomics and Genomics (IPPG), Ludwig-Maximilians-University, Munich, Germany.; Department of Psychiatry and Mental Health, University of Cape Town, Cape Town, South Africa.; The Ernest J. Del Monte Institute for Neuroscience, Department of Neuroscience, University of Rochester School of Medicine and Dentistry.; Neuroscience Research Australia, Sydney, NSW, Australia.; School of Medical Sciences, University of New South Wales, Sydney, New South Wales, Australia.; Department of Psychiatry, University of Vermont, Burlington, VT, USA.; Laboratory of Neuroimaging, Department of Neurology, University of Campinas, Campinas, Brazil.; Department of Psychology, Stanford University, USA.; Department of Psychiatry, Academic Medical Center, University of Amsterdam, Amsterdam, The Netherlands.; Arkin Mental Health & Amsterdam Institute for Addiction Research, Academic Medical Center, University of Amsterdam.; Department of Psychiatry and Psychotherapy, University Medicine Greifswald, Germany.; German Center for Neurodegenerative Diseases (DZNE), Rostock/Greifswald, Germany.; Interdisciplinary Center Psychopathology and Emotion Regulation, University Medical Center Groningen, University of Groningen, The Netherlands.; Section for Experimental Psychopathology and Neuroimaging, Department of General Psychiatry, Heidelberg University, Heidelberg, Germany.; Department of Psychiatry, Haukeland University Hospital, Bergen, Norway.; Queensland Brain Institute, University of Queensland, Brisbane, Australia.; Division of Psychiatry, University of Edinburgh, Edinburgh, UK.; Department of Psychiatry, University Medical Center Groningen, University of Groningen, The Netherlands.; Department of Cognitive Psychology, VU University Amsterdam, Amsterdam, The Netherlands.; Department of Clinical Neuropsychology, VU University Amsterdam, Amsterdam, The Netherlands.; School of Psychological Sciences, University of Melbourne, Melbourne, Australia.; Imaging Genetics Center, Mark and Mary Stevens Neuroimaging and Informatics Institute, Keck School of Medicine of USC, Marina del Rey, CA 90292 USA.; Department of Psychiatry, University of Iowa College of Medicine, Iowa City, Iowa, USA.; Department of Psychology, Stanford University, Stanford, CA, USA.; University of Groningen, University Medical Center Groningen, Department of Psychiatry, Groningen, The Netherlands.; Amsterdam Institute for Addiction Research, Academic Medical Center, University of Amsterdam, Amsterdam, The Netherlands.; Department of Psychiatry, Amsterdam, The Netherlands.; Department of Clinical Medicine, University of Bergen, Bergen, Norway.; Department of Biological and Medical Psychology, University of Bergen, Bergen, Norway.; De Bascule, Academic Center for Child and Adolescent Psychiatry, Amsterdam, The Netherlands.; AMC, Department of Child and Adolescent Psychiatry, Amsterdam, The Netherlands.; Department of Psychiatry, Oregon Health & Science University, Portland, OR, USA.; University Department of Psychiatry, Warneford Hospital, Oxford, UK.; Departments of Cognitive Science, Psychiatry, Radiology, University of California, San Diego, CA, USA.; Center for Human Development, University of California, San Diego, CA, USA.; Norwegian Centre for Mental Disorders Research (NORMENT), K.G. Jebsen Centre for Psychosis Research, Division of Mental Health and Addiction, Oslo University Hospital, Oslo, Norway.; Mental Health Research Center, Moscow, Russia.; Department of Psychiatry, University of Dublin, Trinity College Dublin, Dublin, Ireland.; The Child Study Center at NYU Langone Medical Center, New York, USA.; School of Psychology, Trinity College, Dublin, Ireland.; Trinity College Institute of Neuroscience, Dublin, Ireland.; Clinical NeuroImaging Research Core, National Institute on Alcohol Abuse and Alcoholism, National Institutes of Health, Baltimore, MD, USA.; Beth Israel Deaconess Medical Center, Boston, MA, USA.; Olin Neuropsychiatry Research Center, Hartford CT, USA.; Department of Psychiatry and Psychotherapy, Philipps-University Marburg, Germany.; Child Neuropsychology Section, Department of Child and Adolescent Psychiatry, University Hospital Aachen, Aachen, Germany.; Department of Psychiatry, Washington University School of Medicine, St Louis, MO, USA.; Department of Psychiatry, Seoul National University College of Medicine, Seoul, Republic of Korea.; Department of Brain and Cognitive Sciences, Seoul National University College of Natural Sciences, Seoul, Republic of Korea.; Institute of Neuroscience and Physiology, Sahlgrenska Academy at Gothenburg University, Gothenburg, Sweden.; Department of Child and Adolescent Psychiatry and Psychology, Hospital Clinic, Barcelona, Spain.; August Pi I Sunyer Biomedical Research Institute (IDIBAPS), Barcelona, Spain.; Department of Medicine, University of Barcelona, Barcelona, Spain.; CIBERSAM.; School of Psychiatry, University of New South Wales, Sydney, NSW, Australia.; University of New Mexico, Albuquerque, New Mexico.; Division of Molecular Psychiatry, Center of Mental Health, University of Wurzburg, Wurzburg, Germany.; Laboratory of Psychiatric Neurobiology, Institute of Molecular Medicine, I.M. Sechenov First Moscow State Medical University, Moscow, Russia.; Department of Translational Neuroscience, School for Mental Health and Neuroscience (MHeNS), Maastricht University, Maastricht, The Netherlands.; Beijing Key Laboratory of Applied Experimental Psychology, National Demonstration Center for Experimental Psychology Education (Beijing Normal University), Faculty of Psychology, Beijing Normal University, Beijing, China.; Department of Psychiatry, University of Minnesota, Minneapolis, MN, USA.; SU/UCT MRC Unit on Risk and Resilience in Mental Disorders, Department of Psychiatry, Stellenbosch University, South Africa.; Department of Psychiatry and Biobehavioral Sciences, University of California, Los Angeles, CA, USA.; Institute of Psychology Health and Society, University of Liverpool, Liverpool, UK.; Behavioural Science Institute, Radboud University, Nijmegen, The Netherlands.; Departments of Psychiatry and Pediatrics, University of Calgary, Calgary AB, Canada.; Child and Adolescent Imaging Research Program, Alberta Children's Hospital, Calgary AB, Canada.; Mathison Centre for Mental Health Research & Education, Hotchkiss Brain Institute, University of Calgary, Calgary AB, Canada.; Strategic Clinical Network for Addictions and Mental Health, Alberta Health Services, Calgary AB, Canada.; Alberta Children's Hospital Research Institute, Calgary AB, Canada.; Institut des Maladies Neurodégénératives, UMR5293. Groupe d’Imagerie Neurofonctionnelle, CEA - CNRS - Université de Bordeaux, Bordeaux, France.; Developmental Imaging, Murdoch Childrens Research Institute, Royal Children's Hospital, Melbourne, Australia.; Department of Research and Education, Oslo University Hospital, Oslo, Norway.; Institute of Clinical Medicine, University of Oslo, Oslo, Norway.; Department of Psychiatry and Psychology, University of Barcelona, Barcelona, Spain.; Department of Psychiatry, Trinity College Dublin, Dublin, Ireland.; Black Dog Institute, Prince of Wales Hospital, Randwick, NSW, Australia.; Prince of Wales Hospital, Sydney, NSW, Australia.; Institut des Maladies Neurodégénératives, UMR 5293 Groupe d’Imagerie Neurofonctionnelle, CEA - CNRS - Université de Bordeaux.; Department of Neuroscience, Cumming School of Medicine, University of Calgary, Calgary, AB, Canada.; Alberta Children's Hospital Research Institute, Calgary, AB, Canada.; Centre for Advanced Imaging, University of Queensland, Brisbane, Australia.; Neuropsychopharmacology Unit, Division of Brian Sciences, Imperial College London, London, UK.; Sunnaas Rehabilitation Hospital HT, Nesodden, Norway.; Center for MR-Research, University Children's Hospital, Zurich, Switzerland.; Emma Children’s Hospital Amsterdam Medical Center, Amsterdam, The Netherlands.; VU Medical Center, Amsterdam, The Netherlands.; Clinical Neuropsychology section, Vrije Universiteit Amsterdam, Amsterdam, The Netherlands.; Laureate Institute for Brain Research, Tulsa, Oklahoma, USA.; Department of Psychiatry, University of California San Diego, La Jolla, California, USA.; Child and Adolescent Mental Health Center, Capital Region, Denmark.; Centre for Psychiatric Research and Education, Department of Clinical Neuroscience, Karolinska Institutet, Stockholm, Sweden.; Department of Psychosis Studies, Institute of Psychiatry, Psychology, and Neuroscience, King's College London, UK.; Department of Psychiatry and Legal Medicine, Universitat Autonoma de Barcelona, Barcelona, Spain.; Department of Psychiatry, Hospital Universitari Vall d’Hebron, CIBERSAM, Barcelona, Spain.; Department of Psychiatry, Psychosomatic Medicine and Psychotherapy, University Hospital Frankfurt, Frankfurt, Germany.; Department of Psychiatry and Behavioral Sciences, Stanford University, USA.; Neuropsychiatric Institute, Prince of Wales Hospital, Randwick, Australia.; Centre for Youth Mental Health, The University of Melbourne, Melbourne, Australia.; Department of Child and Adolescent Psychiatry, University Hospital Aachen, Aachen, Germany.; Neurobehavioral Clinical Research Section, National Human Genome Research Institute, Bethesda, USA.; National Institute of Mental Health, Bethesda, MD, USA.; Department of Paediatrics, University of Melbourne, Melbourne, Australia.; Department of Psychiatry, University of California, San Diego, 9500 Gilman Dr., La Jolla, CA, USA.; Veterans Affairs San Diego Health Care System, La Jolla, CA, USA.; Max Planck Institute for Human Cognitive and Brain Sciences, Department of Neurology, Leipzig, Germany.; Leiden University, Institute of Psychology, Cognitive Psychology Unit & Leiden Institute for Brain and Cognition, Leiden, The Netherlands.; School of Psychology and Illawarra Health and Medical Research Institute, University of Wollongong, Wollongong, Australia.; Minneapolis VA Health Care System & University of Minnesota, Minneapolis, MN, USA.; SU/UCT MRC Unit on Risk and Resilience in Mental Disorders, Department of Psychiatry and Mental Health, University of Cape Town, South Africa.; Clinical Neuroscience and Development Laboratory, Olin Neuropsychiatry Research Center, Hartford CT, USA.; Child & Adolescent Research, Hartford Hospital/The Institute of Living, Hartford CT, USA.; Department of Psychiatry, Yale University School of Medicine, Hartford CT, USA.; National laboratory of Pattern Recognition, Institute of Automation, Chinese Academy of Sciences, Beijing, China.; Department of Pediatrics, Division of Behavioral Medicine and Clinical Psychology, Cincinnati Children's Hospital Medical Center, Cincinnati, OH, USA.; Department of Psychiatry and Behavioral Sciences, University of New Mexico, Albuquerque, NM, USA.; Department of Psychiatry, University of New Mexico, Albuquerque, NM, USA.; Department of Psychology and Neuroscience Institute, Georgia State University, Atlanta GA 30302.; Department of Psychiatry and Human Behavior, University of California, Irvine, Irvine, USA.; K.G. Jebsen Centre for Psychosis Research / Norwegian Centre for Mental Disorder Research (NORMENT), Institute of Clinical Medicine, University of Oslo, Oslo, Norway.; School of Psychology, University of Queensland, Brisbane, Australia.; Academic Child Psychiatry Unit, Royal Children's Hospital, University of Melbourne, Melbourne, Victoria, Australia.; Charite - Universitatsmedizin Berlin, corporate member of Freie Universitat Berlin, Humboldt-Universitat zu Berlin, and Berlin Institute of Health, Department of Psychiatry and Psychotherapy, Campus Mitte, Berlin, Germany.; Fundació IMIM, Barcelona, Spain.; INNDACYT, Barcelona, Spain.; Department of Psychiatry, University of Toronto, Toronto, Canada.; Institute for Community Medicine, University Medicine Greifswald, Germany.; DZHK (German Centre for Cardiovascular Research), partner site Greifswald, Germany.; German Centre for Diabetes Research (DZD), Site Greifswald, Germany.; Department of Psychology, Georgia State University, Atlanta GA, USA.; Brain Research Imaging Centre, Centre for Clinical Brain Sciences and Dementia Research Institute at the University of Edinburgh, Edinburgh, UK.; Scottish Imaging Network, A Platform for Scientific Excellence (SINAPSE) Collaboration, Edinburgh, UK.; Centre for Clinical Brain Sciences, Centre for Cognitive Ageing and Cognitive Epidemiology, and UK Dementia Research Institute at The University of Edinburgh, Edinburgh, UK.; NORMENT, KG Jebsen Centre for Psychosis Research, Division of Mental Health and Addiction, Oslo University Hospital & Institute of Clinical Medicine, University of Oslo, Oslo, Norway.; Department of Psychology, University of Oslo, Oslo, Norway.; Department of Child and Adolescent Psychiatry, Erasmus University Medical Centre, Rotterdam, Netherlands.; Department of Radiology, Erasmus University Medical Centre, Rotterdam, Netherlands.; Monash Institute of Cognitive and Clinical Neurosciences and School of Psychological Sciences, Monash University, Melbourne, Australia.; Seoul National University Hospital, Seoul, Republic of Korea.; Yeongeon Student Support Center, Seoul National University College of Medicine, Seoul, Republic of Korea.; Institute of Health and Biomedical Innovation, Queensland University of Technology, Brisbane, Australia.; Members of Karolinska Schizophrenia Project (KaSP) are listed before References.; Department of Psychiatry, Yale University, New Haven, CT, USA.; Olin Neuropsychiatric Research Center, Hartford, CT, USA.; Department of Human Genetics, Donders Institute for Brain, Cognition and Behaviour, Radboud University Medical Center, Nijmegen, The Netherlands.; Department of Psychiatry, Donders Institute for Brain, Cognition and Behaviour, Radboud University Medical Center, Nijmegen, The Netherlands.; Donders Institute for Brain, Cognition and Behavior, Radboud University, Nijmegen, The Netherlands.; Centre for Psychiatry Research, Department of Clinical Neuroscience, Karolinska Institutet, & Stockholm County Council, Stockholm, Sweden; Department of Physiology and Pharmacology, Karolinska Institutet, Stockholm, Sweden; Neuroimmunology Unit, Department of Clinical Neuroscience, Karolinska Institutet, Stockholm, Sweden

**Author notes:** **Correspondence Author:** Clyde Francks, Ph.D., Xiang-Zhen Kong, Ph.D.

**Keywords:** brain asymmetry, lateralization, cortical thickness, surface area, meta-analysis

## Abstract

Hemispheric asymmetry is a cardinal feature of human brain organization. Altered brain asymmetry has also been linked to some cognitive and neuropsychiatric disorders. Here the ENIGMA consortium presents the largest ever analysis of cerebral cortical asymmetry and its variability across individuals. Cortical thickness and surface area were assessed in MRI scans of 17,141 healthy individuals from 99 datasets worldwide. Results revealed widespread asymmetries at both hemispheric and regional levels, with a generally thicker cortex but smaller surface area in the left hemisphere relative to the right. Regionally, asymmetries of cortical thickness and/or surface area were found in the inferior frontal gyrus, transverse temporal gyrus, parahippocampal gyrus, and entorhinal cortex. These regions are involved in lateralized functions, including language and visuospatial processing. In addition to population-level asymmetries, variability in brain asymmetry was related to sex, age, and brain size (indexed by intracranial volume). Interestingly, we did not find significant associations between asymmetries and handedness. Finally, with two independent pedigree datasets (*N* = 1,443 and 1,113, respectively), we found several asymmetries showing modest but highly reliable heritability. The structural asymmetries identified, and their variabilities and heritability provide a reference resource for future studies on the genetic basis of brain asymmetry and altered laterality in cognitive, neurological, and psychiatric disorders.

**Significance Statement:** Left-right asymmetry is a key feature of the human brain's structure and function. It remains unclear which cortical regions are asymmetrical on average in the population, and how biological factors such as age, sex and genetic variation affect these asymmetries. Here we describe by far the largest ever study of cerebral cortical brain asymmetry, based on data from 17,141 participants. We found a global anterior-posterior 'torque' pattern in cortical thickness, together with various regional asymmetries at the population level, which have not been previously described, as well as effects of age, sex, and heritability estimates. From these data, we have created an on-line resource that will serve future studies of human brain anatomy in health and disease.

---

Understanding the functional specialization of the cerebral hemispheres is a long-standing and central issue in human neuroscience research. At the population-level, hemispheric asymmetry, or lateralization, is involved in various perceptual and cognitive functions, including language (1, 2), face processing (3–5), visuospatial processing (3, 6, 7), and reasoning (8, 9), as well as handedness (10). For example, language lateralization involves leftward dominance for various processes involved in speech perception and production in most people (1, 2). Moreover, altered hemispheric lateralization has been associated with numerous cognitive and neuropsychiatric disorders, including dyslexia (11), Alzheimer’s disease (12), attention-deficit/hyperactivity disorder (ADHD) (13), psychotic disorders (14–17), autism (18) and mood disorders (19, 20). Various aspects of brain asymmetry, including anatomical asymmetries of perisylvian language-related cortical regions, appear *in utero* during the second trimester of gestation (21, 22). Thus, brain laterality is likely to be under the control of genetic-developmental programs which are inherently lateralized, such as those that have been described for the left-right visceral axis (affecting the placement of the heart, lungs etc.) (23, 24). Together, these observations indicate that asymmetry is a core element of the brain’s usual organization, which is required for optimal functioning and influenced by genetic factors.

Although structural and functional asymmetries are likely to be interrelated in the typically lateralized human brain, the nature of structure-function relations are far from clear. For example, it is still not understood whether anatomical asymmetries around the Sylvian fissure are an important aspect of left-hemisphere language dominance (25, 26). Furthermore, variations in structural and functional asymmetry have been reported to correlate poorly (27–30), which further complicates assessment of the structure-function relations and dependencies. The literature has, however, been based on generally small sample sizes and heterogeneous methods for assessing asymmetries and their variabilities (31), leading to confusion about which structures are actually anatomically asymmetrical at the population level, and to what degrees (see below). This has also been the case for asymmetry-disorder studies. In this context, and as motivation for the present study, it is important to characterize anatomical asymmetries in a large sample of healthy individuals, in order to provide a definitive and normative reference for future studies of hemispheric specialization in both healthy and clinical populations.

One aspect of structural asymmetry in the human brain is “Yakovlevian torque”, an overall hemispheric twist giving rise to the frontal and occipital petalia, which describes protrusions of the right frontal and left occipital regions over the midline (32–34). At a regional level, later studies that applied computational methods to MRI data mainly focused on volumetric measures of cortical structures and revealed both replicable and inconsistent findings of asymmetries. For example, Goldberg et al. (2013) summarized in their study that regions implicated in visual processing show rightward volumetric asymmetries, while, in contrast, somatosensory, auditory, and parts of the premotor cortices show leftward volumetric asymmetries (35). One recent study replicated this distribution of regional asymmetries, especially in the lateral view (36), but several studies have shown quite different asymmetry results (33, 37, 38). For example, Goldberg et al. (2013) and Esteves et al. (2017) found a greater superior frontal volume in the left hemisphere, while Watkins et al. (2001) found greater superior frontal volume in the right.

Cortical volume is, by definition, a product of two distinct aspects of the brain, i.e. cortical thickness and surface area (39, 40); researchers have also attempted to assess the asymmetries of cortical thickness and surface area separately, using neuroimaging surface-based approaches (41, 42). Regarding cortical thickness, a number of studies have found mixed results for asymmetry patterns. For example, Luders et al. (2006) found greater left-sided thickness in parts of the cingulate, precentral gyrus, orbital frontal gyrus, and temporal and parietal lobes, and greater right-sided thickness in the inferior frontal gyrus (43). However, other studies (44–47) revealed somewhat inconsistent patterns of thickness asymmetry. For instance, Zhou et al. (2013), studying individuals of an age range similar to that in Luders et al. (2006), did not find leftward asymmetry in the precentral gyrus, but revealed a strong rightward asymmetry in the lateral parietal and occipital regions. For an overview of mixed results of asymmetry patterns observed in previous studies, please refer to Figure S1 in Supplemental Information. Regarding regional surface area asymmetries, some repeatable findings have been found for the supramarginal gyrus (leftward) (44, 45, 48), the middle temporal gyrus (rightward) (44, 45),
and the anterior cingulate gyrus (rostral: leftward; caudal: rightward) (44, 45). However, there are also many inconsistent results across studies, such as for the lateral occipital cortex, which showed a strong rightward asymmetry in Chiarello et al. (2016) (49), but leftward asymmetry in Koelkebeck et al. (2016) (see a summary in (50)). These mixed results of brain structural asymmetry may reflect differences in many factors, including statistical power and confidence intervals related to sample sizes as well as differences in scanning, brain segmentation, and parcellation methods. Thus, a large-scale survey using harmonized approaches is needed to give a clearer picture of the lateralization in the human brain.

Another potential source for the mixed results in the literature is variability across individuals and in relation to factors like age and sex (51–53). For example, a recent study has observed that males show, on average, more pronounced gray matter volume asymmetries in superior temporal language regions than females (54). Changes in structural asymmetries with age have also been reported (13, 55), but not consistently (47). Another potential factor linked to brain lateralization is handedness, although the associations are very weak as reported (30, 45, 56). For example, with more than 100 left-handed participants and roughly 2000 right-handed participants, Guadalupe et al. (2014) suggested an association of handedness with the surface area of the left precentral sulcus, but this was not significant after multiple testing adjustment. In addition, greater cortical asymmetry has been observed in participants with larger overall brain size (50). Thus, the existing literature on variability in brain structural asymmetries suggests influences of individual differences in age, sex, handedness, and brain size, but again a large-scale study is needed to clarify the nature of any such relations. The largest previous studies of brain asymmetries were conducted by Plessen et al. (2014) and Zhou et al. (2013) in relation to sex and age in sample sizes of 215 and 274 participants, and Maingault et al. (2016) in a sample size of 250 (120 left-handers) in relation to handedness. Each of these studies used different methodological approaches. Thus, a large-scale study of thousands of participants would be a major step forward in achieving a more accurate description of the typical asymmetries of the human brain, as well as variation in these asymmetries and some key biological factors which affect them.

The ENIGMA consortium provides the opportunity for large-scale meta-analysis studies of brain anatomy based on tens of thousands of participants with structural MRI data (57). In this study, we present the largest analysis of structural asymmetries in the human cerebral cortex, with MRI scans of 17,141 healthy individuals from 99 datasets worldwide, in a harmonized multi-site study using meta-analytic methods. Our aim was to identify cortical regions that consistently show asymmetry with regard to either cortical thickness or surface area, to provide a clear picture of population-level asymmetries in the human brain. We also assessed potential influences of age, sex, handedness, and brain size (indexed by intracranial volume, ICV) on the variability in asymmetries, as well as of the methodological factor of MRI scanner field strength. Furthermore, as a first step towards elucidating the genetic basis of variability in brain asymmetry, we further analyzed two independent pedigree datasets, i.e. the Genetics of Brain Structure (GOBS; N = 1,443) and Human Connectome Project (HCP; N = 1,113) datasets, to estimate heritability of the asymmetry measures.

## Results

### Meta-analysis of population-level asymmetry

Meta-analysis of population-level asymmetry revealed widely distributed asymmetries in both cortical thickness and surface area. Specifically, we found global differences between the two hemispheres, with generally thicker cortex in the left hemisphere (*b* = 0.13, *Z* = 3.64, *p* = 0.00040; Figure 2), but larger surface area in the right hemisphere (*b* = −0.33, *Z* = −11.30, *p* = 1.36e-29; Figure 3).

**Figure. 2.**
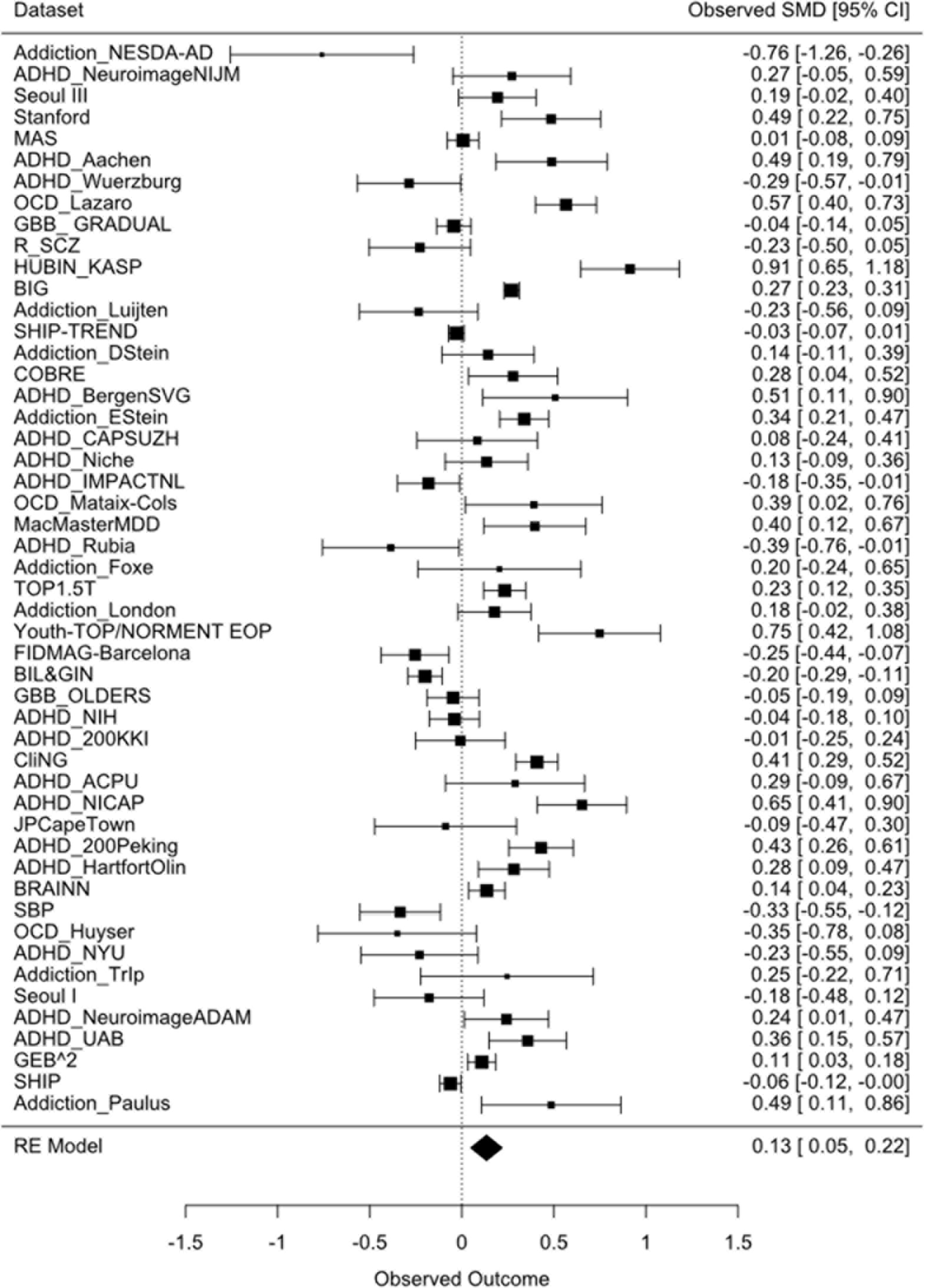
Forest plot of asymmetry score per dataset, for the overall asymmetry in cortical thickness. Asymmetry score indicates the effect size of the inter-hemispheric difference. The size of a square is proportional to the weights assigned in meta-analysis. The confidence intervals are shown, as well as a dashed vertical line to indicate the point of an asymmetry score of zero. The forest plot was based 50 randomly selected datasets for visualization. Results based on all datasets can be found via http://conxz.github.io/neurohemi/.

**Figure. 3.**
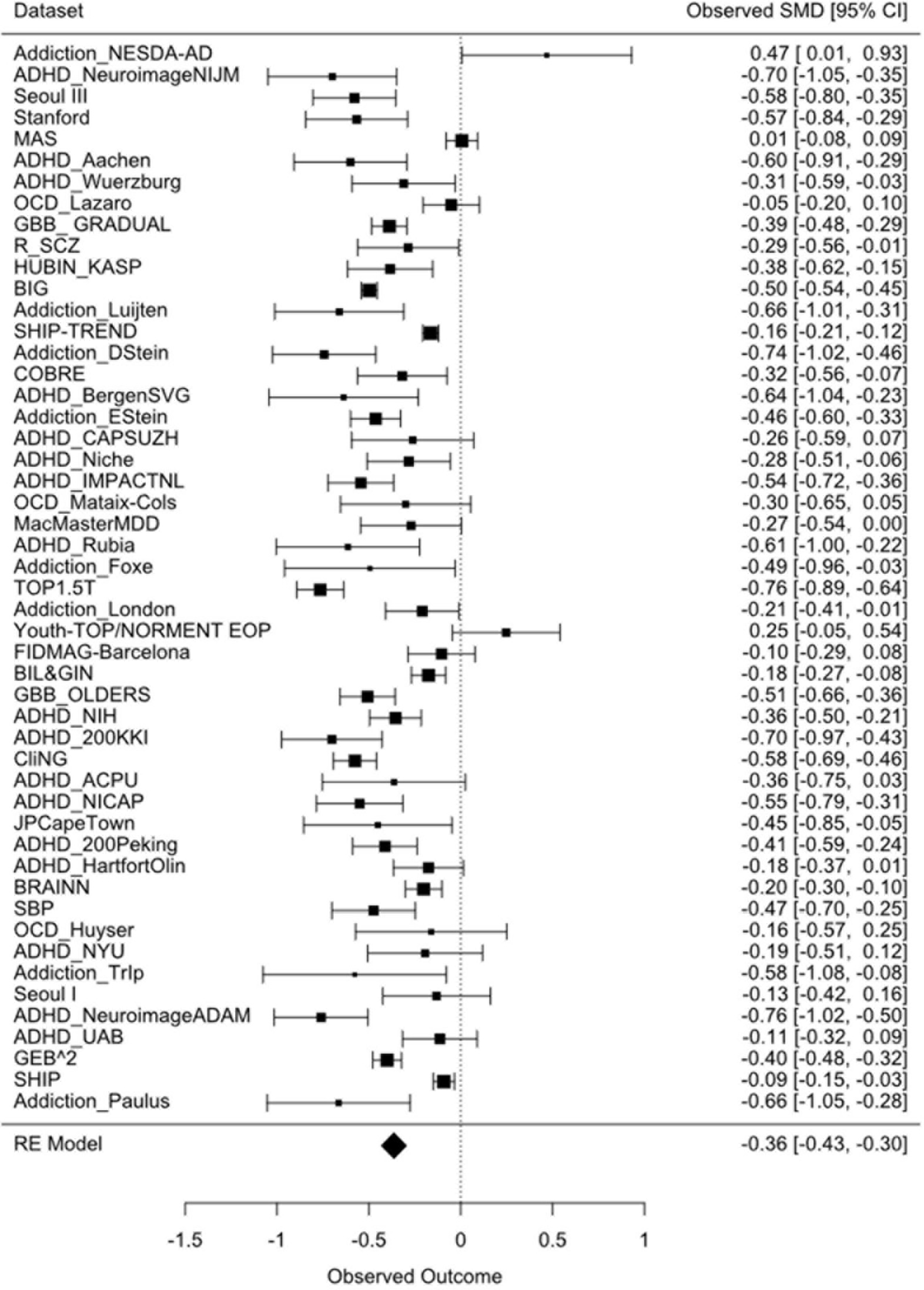
Forest plot of asymmetry score per dataset, for the overall asymmetry in surface area. Asymmetry score indicates the effect size of the inter-hemispheric difference. The size of a square is proportional to the weights assigned in meta-analysis. The confidence intervals are shown, as well as a dashed vertical line to indicate the point of an asymmetry score of zero. The forest plot was be 50 randomly selected datasets for visualization. Results based on all datasets can be found via http://conxz.github.io/neurohemi/.

Substantial, regionally specific differences between the two hemispheres were also observed for both cortical thickness and surface area. In terms of cortical thickness, 76.5% (26/34) of the regions showed significant asymmetry, after correcting for multiple comparisons (*p* <0.05, Bonferroni corrected). Specifically, regions showing significant leftward asymmetry (i.e., left > right) of cortical thickness were identified in the anterior cortex, including the lateral, dorsal and medial frontal cortex, the primary sensory, superior parietal, cingulate, and medial temporal cortices (Fig. 4; see Supplemental Information S3). In contrast, rightward asymmetry (i.e., right > left) was prominent in the posterior regions, including lateral and medial parts of the temporal, parietal, and occipital cortices. This fronto-occipital asymmetry pattern in cortical thickness is striking (Fig. 4) and may also relate to the petalia and Yakovlevian torque effects described above (also see Discussion below). In addition, three temporal regions (especially the inferior temporal and fusiform gyri) showed a trend of rightward asymmetry as defined by uncorrected P <0.05 (inferior temporal: b = −0.11, Z = −2.92, uncorrected p = 0.0035; fusiform: b = −0.09, Z = −2.64, uncorrected p = 0.0082; middle temporal: b = −0.10, Z = −2.19, uncorrected p = 0.029).

**Figure. 4.**
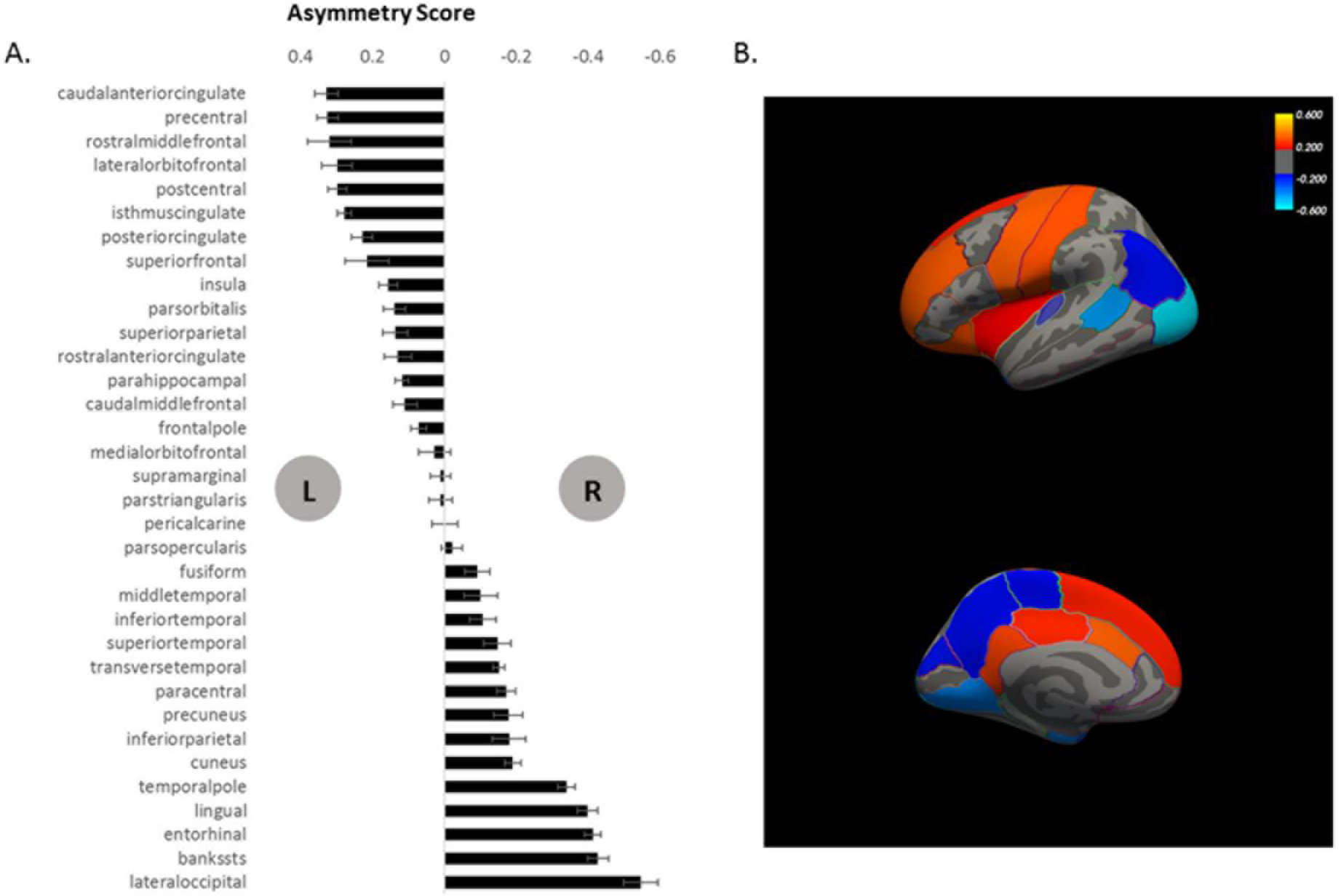
Average regional asymmetries in cortical thickness reveal a fronto-occipital pattern. Positive asymmetry (left side in A; red in B) indicates leftward asymmetry, while negative asymmetry (right side in A; blue in B) indicates rightward asymmetry. Asymmetry score indicates the effect size of the inter-hemispheric difference. Error bars indicate standard error of the mean. L, left; R, right.

Similarly, 91.1% (31/34) of the regions showed significant asymmetries of their surface areas after correcting for multiple comparisons (*p* <0.05, Bonferroni corrected). However, unlike for thicknesses, the surface area asymmetries showed no obvious leftward or rightward patterns involving multiple neighboring areas, or generally along the fronto-occipital axis (Fig. 5; see Supplemental Information S3). Two language-related regions showed the largest leftward asymmetries of surface area, which were the opercular part of the inferior frontal gyrus (posterior part of the Broca's area) and the transverse temporal gyri (Heschl's gyri). In contrast, however, another two language-related regions, i.e. the triangular part of the inferior frontal gyrus (anterior part of the Broca’s area) and the inferior parietal gyrus, showed strong rightward asymmetries of surface area. These findings suggest that opposite asymmetries in morphology of regions within a given network (i.e., language network), or within one functional area (the Broca’s area), might be linked to different roles of each constituent part (see Discussion below).

**Figure. 5.**
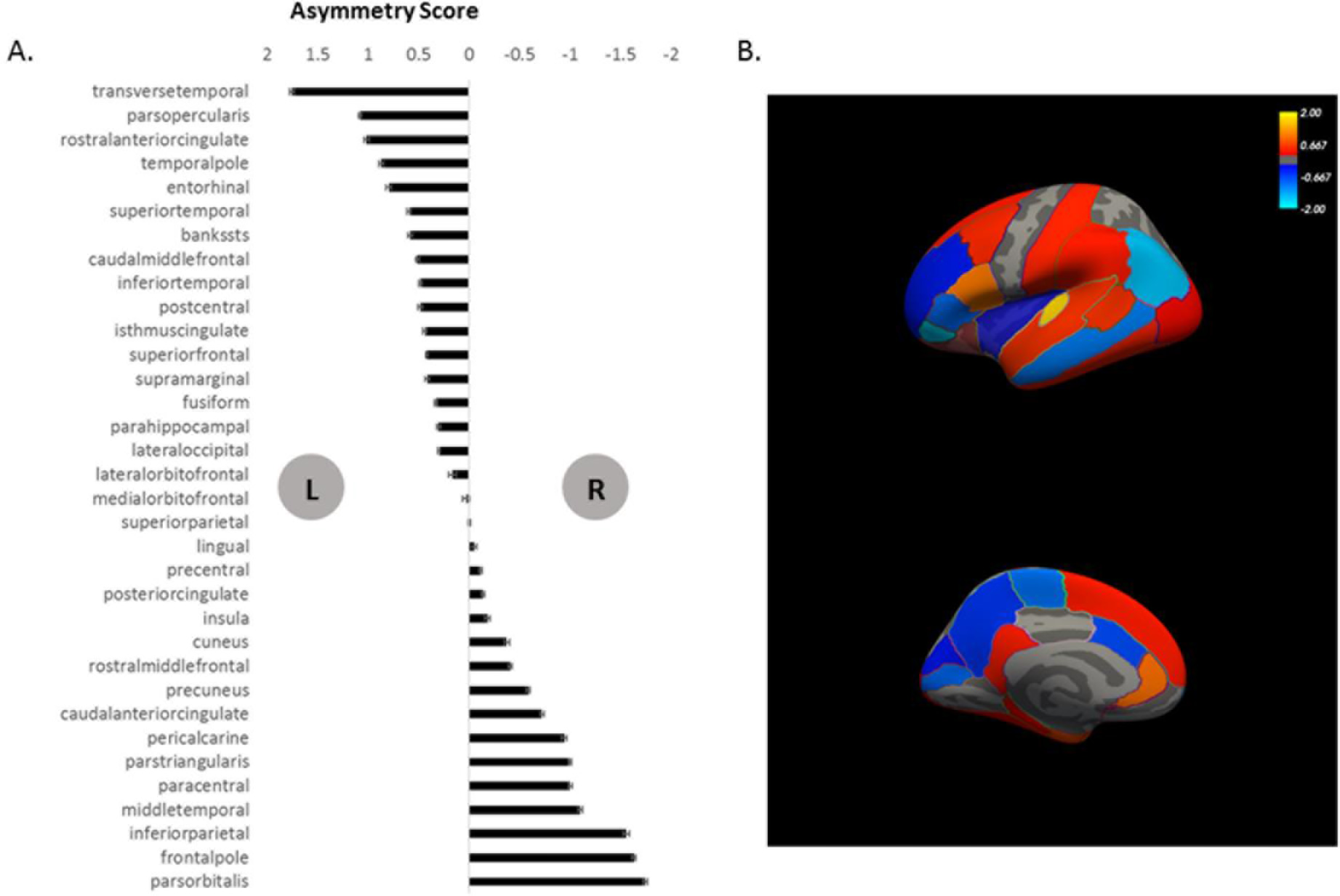
Average asymmetry pattern in surface area. Positive asymmetry (left side in A; red in B) indicates leftward asymmetry, while negative asymmetry (right side in A; blue in B) indicates rightward asymmetry. Asymmetry score indicates the effect size of the inter-hemispheric difference. Error bars indicate standard error of the mean. L, left; R, right.

Effect sizes of cortical thickness and surface area were found to be independent, as illustrated by the absence of a significant correlation between thickness and surface area asymmetries across all cortical regions (*r* = −0.14, *p* = 0.416).

### Moderator analyses using meta-regression

As shown above (e.g., Fig. 2, 3 and Supplemental Information S3), we observed moderate to substantial heterogeneity in the asymmetry distributions across datasets (I^2^ ranges from 36% to 98%).

To further address the heterogeneity across the samples included in the meta-analyses, we investigated several moderating variables, including sex ratio, median age, handedness ratio, and median ICV. Moderator analyses revealed an influence of the median age of samples on the global hemispheric difference in surface area (*Z* = 2.09, *p* = 0.036), suggesting a reduced rightward asymmetry with increasing age. No other potential moderators showed significant effects on global cortical thickness or surface area asymmetry (*p* > 0.10). Moderator analyses for each specific region suggested an influence of the median age of samples on the asymmetry of the surface area of the paracentral gyrus (*Z* = −4.35, *p* = 1.38e-5), and an influence of median ICV on the asymmetry of the surface area in the insula (*Z* = −3.18, *p* = 0.0014). Given that both the paracentral gyrus and insula showed significant rightward asymmetry in surface area, these findings indicate a decreasing rightward asymmetry with increasing age and with brain size, respectively. In addition, we found a significant effect of scanner field strength on the surface area asymmetry in the insula (*Z* = 4.12, *p* = 3.82e-5). However, this could be largely reduced by including the other moderating variables (i.e., sex ratio, median age, handedness ratio, and median ICV) in the analysis (*Z* = 2.35, *p* = 0.019).

### Meta-analysis of sex effects on cortical asymmetries

No significant sex effect on the global asymmetry of cortical thickness was found (p > 0.10), but notable regionally specific effects on thickness asymmetries were observed in the medial temporal regions (Figure 6), including the parahippocampal gyrus (*Z* = 3.57, *p* = 0.00036) and the entorhinal cortex (*Z* = 3.61, *p* = 0.00030), after correcting for multiple comparisons. Together with the population-level asymmetry observed above, these results indicate that males show more leftward and less rightward asymmetry in cortical thickness of the parahippocampal gyrus and the entorhinal cortex, respectively.

We found a significant sex difference in global asymmetry of surface area (*Z* = −2.62, *p* = 0.0088), indicating that males have more rightward overall asymmetry in surface area, compared with females. In addition, meta-regression analysis showed that this effect changed with the median ages of samples: we found larger effects of sex (females > males) in the younger samples, compared to the older samples (*Z* = 2.80, *p* = 0.0052). Regionally specific effects of sex on surface area asymmetry were also revealed (Figure 6; Bonferroni corrected), located in the frontal (superior frontal gyrus, the pars orbitalis region of the left inferior frontal gyrus), temporal (superior temporal gyrus, temporal pole, parahippocampal gyrus and fusiform gyrus), parietal (inferior parietal gyrus and supramarginal gyrus), and anterior cingulate cortices. In addition, various other regions showed nominally significant sex effects (uncorrected p < 0.05) without surviving correction for multiple comparisons. More information can be seen in Supplemental Information S4.

We did not observe a significant correlation across regions of the sex effects on the asymmetries of cortical thickness and surface area (*r* = 0.14, *p* = 0.434), further supporting the independent nature of these two cortical features.

**Figure. 6.**
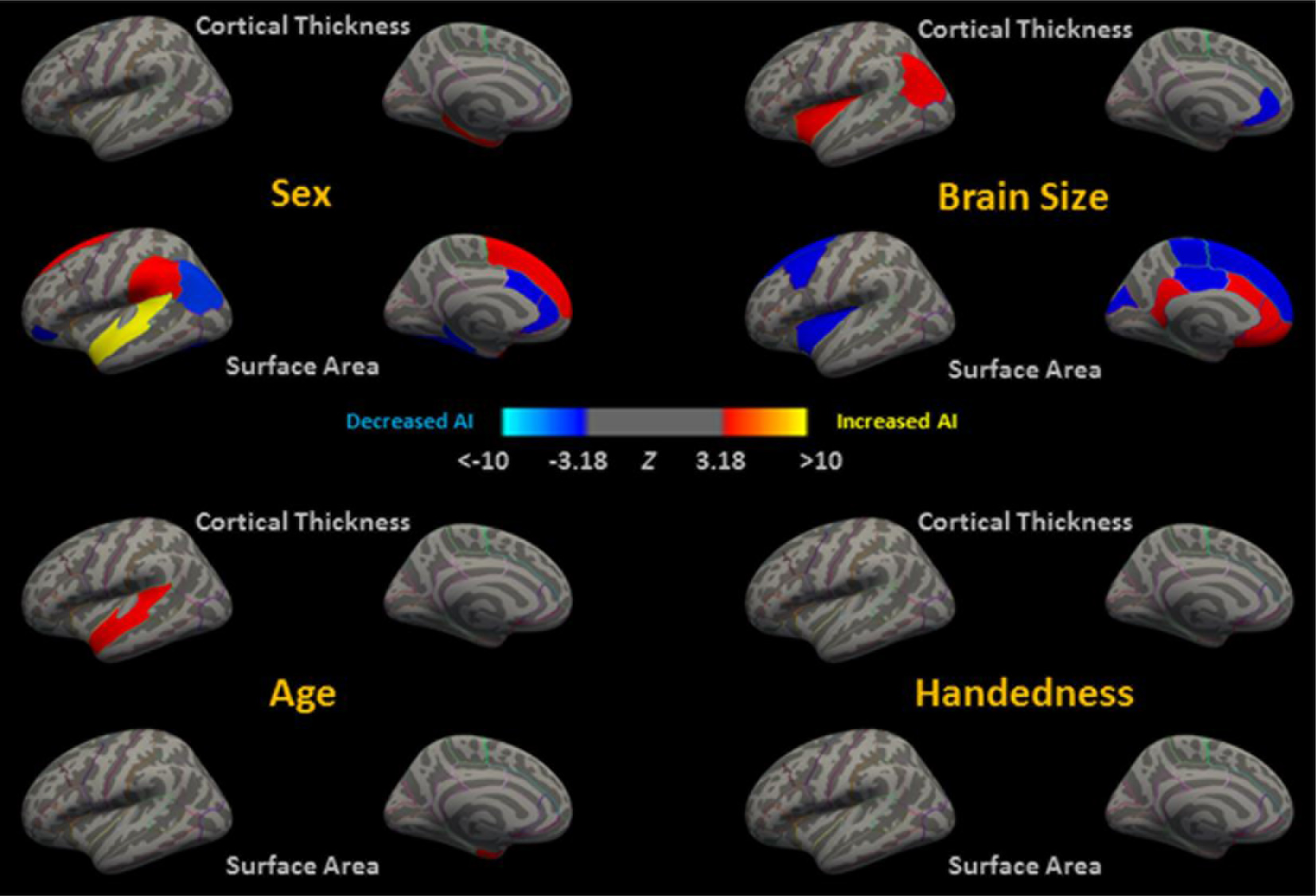
Meta-analysis results for effects of sex, age, brain size (indexed by ICV), and handedness on regional asymmetry indexes in cortical thickness and surface area. Red-yellow indicates an increased asymmetry index (AI) in males/with age and brain size; blue-lightblue indicates a decreased AI in males/with age and brain size. AI was defined as (L-R)/((L+R)/2). A *Z* threshold of 3.18 (*p* = 0.05, Bonferroni corrected) was used. For more details, please see Supplemental Information S8.

### Meta-analysis of age effects on cortical asymmetries

An initial analysis of samples with an age range of larger than five years showed no significant effects of age on global asymmetries of either cortical thickness or surface area (*ps* > 0.10). Several regionally specific, nominally significant effects were found: the superior temporal gyrus (cortical thickness: *Z* =2.38, *p* =0.017), the banks of superior temporal sulcus (surface area: *Z* = −1.97, *p* = 0.049), and the entorhinal cortex (surface area: *Z* = 2.84, *p* = 0.0045). However, when restricting the analysis to only those datasets with wider age ranges (at least 20 years range), we observed significant age effects. Specifically, increasing age was associated with more pronounced leftward overall asymmetry in cortical thickness (*Z* = 2.65, *p* = 0.0081), which partly reflects a similar age effect on the thickness asymmetry of the superior temporal gyrus (*Z* = 3.17, *p* = 0.0015; Figure 6). In addition, a similar effect on regional surface area asymmetry was observed in the entorhinal cortex (Z = 3.21, p = 0.0013). An age effect on surface area asymmetry of the banks of the superior temporal sulcus was nominally significant (*Z* = −1.96, *p* = 0.050). More information can be found in Supplemental Information S5. No significant correlation was found between the age effects across regions on cortical thickness asymmetry and surface area asymmetry (*p*s > 0.05 for both age-range thresholds).

### Meta-analysis of group differences by handedness on cortical asymmetries

We did not find significant associations of handedness with cortical asymmetries, even with this unprecedented sample size (from 555 to 608 left-handers versus 6,222 to 7,243 right-handers, depending on the specific regional asymmetry measure). Some temporal regional surface area asymmetries showed nominally significant associations with handedness, including the fusiform gyrus: *Z* = 2.00, *p* = 0.046; the parahippocampal gyrus: *Z* = −2.33, *p* = 0.020; and the superior temporal gyrus: *Z* = −2.04, *p* = 0.042. More information can be found in Supplemental Information S6. No significant correlation was found between the handedness effects across regions for cortical thickness asymmetry and surface area asymmetry (*r* = −0.15, *p* = 0.403).

### Meta-analysis of ICV effects on cortical asymmetry

ICV showed a significant positive effect (i.e., increased leftward asymmetry) on the overall asymmetry in cortical thickness (*Z* = 2.14, *p* = 0.032). Similar regionally specific effects on cortical thickness asymmetry were found for the inferior parietal gyrus (*Z* = 4.51, *p* = 6.53e-6), and insula (*Z* = 3.71, *p* = 0.00021). A negative effect of greater ICV (i.e., decreased leftward asymmetry) was seen in the rostral anterior cingulate gyrus (*Z* = −5.23, *p* = 1.68e-7) (Figure 6). No significant effect of ICV was found for the overall asymmetry in surface area (*p* > 0.10), but a number of regionally specific effects were revealed (in different directions). Positive effects of greater ICV (i.e., increased leftward/decreased rightward asymmetry) were observed in the medial orbitofrontal gyrus (*Z* = 4.17, *p* =3.10e-5), two anterior cingulate gyri (caudal: *Z* = 5.71, *p* =1.10e-8; rostral: *Z* = 5.67, *p* =1.45e-8), and the isthmus cingulate gyrus (*Z* = 4.32, *p* =1.59e-5) (Figure 6). In addition, negative effects of greater ICV (i.e., increased rightward/decreased leftward asymmetry) were seen for the superior frontal gyrus (*Z* = −6.58, *p* =4.82e-11), the caudal middle frontal gyrus (*Z* =−3.65, *p* =0.00026), the paracentral gyrus (*Z* =−5.19, *p* = 2.11e-7), the insula (*Z* = −5.92, *p* =3.13e-9), the posterior cingulate gyrus (*Z* =−3.24, *p* =0.0012), and the cuneus (*Z* =−4.49, *p* =7.12e-6) (Figure 6). More information can be seen in Supplemental Information S7. Similar to the other factors investigated, no significant correlation was found between the effects of ICV across regions on cortical thickness and surface area asymmetry (*r* = −0.14, *p* = 0.417).

### Heritability of cerebral cortical anatomical asymmetries

In the GOBS dataset, the overall hemispheric asymmetries of both cortical thickness and surface area showed low but statistically significant heritabilities (cortical thickness asymmetry: h^2^ = 0.10, *p* = 0.005; surface area asymmetry: h^2^ = 0.17, *p* = 0.00024). Consistent with a previous study on the genetics of brain structure (58, 59), regional surface area asymmetries seemed to be generally more heritable than thickness asymmetries. The most heritable asymmetries in regional cortical thickness were found in the isthmus (h^2^ = 0.17) and caudal anterior cingulate gyrus (h^2^ = 0.13), the superior (h^2^ = 0.13) and rostral middle frontal gyrus (h^2^ = 0.18), the parahippocampal gyrus (h^2^ = 0.15), and the lateral occipital gyrus (h^2^ = 0.16) (*p* < 0.05, Bonferroni corrected; Table 1). The most heritable asymmetries in regional surface area were found in the entorhinal cortex (h^2^ = 0.24), the superior temporal gyrus (h^2^ = 0.19), the inferior parietal gyrus (h^2^ = 0.19), and the isthmus cingulate gyrus (h^2^ = 0. 17) (*p* < 0.05, Bonferroni corrected; Table 1). For each of these regions, we also estimated the genetic correlation between the measures of the left and right structures. While these correlations were high (indicating high pleiotropy), all were significantly different from 1 (see Table 1). These results indicate that most genetic effects on structural variation are shared bilaterally, but some independent genetic effects exist on each hemisphere, which constitute the heritable contributions to structural asymmetry. Finally, we found that the heritability of most of these regions was validated in the HCP dataset. For more details, please see Supplemental Information S8.

**Table 1.** Significant heritabilities for asymmetry measures based on the GOBS family dataset. In the left part of the table are the heritability and p-values; in the middle part are the genetic correlations between the left and right structural measures, and p-values for whether the genetic correlations differ significantly from 0 or 1. In the right part of the table are the environmental and phenotypic correlation estimates between the left and right regions.

| Metric | Structure | AI heritability |  | Genetic correlation between Left and Right |  |  | Phenotypic (rho-phen) and environmental (rho-env) correlation between Left and Right |  |
| --- | --- | --- | --- | --- | --- | --- | --- | --- |
|  |  | h2 | p-value | rho | p(rho=0) | p(rho=1) | rho-phen | rho-env |
| Cortical Thickness | Hemisphere | 0.102 | 0.0051 | 0.993 | 4.37E-22 | 0.024 | 0.950 | 0.899 |
|  | caudalanteriorcingulate | 0.129 | 0.00037 | 0.718 | 0.0000016 | 0.0023 | 0.285 | 0.113 |
|  | isthmuscingulate | 0.171 | 0.00026 | 0.823 | 4.64E-19 | 0.000045 | 0.546 | 0.270 |
|  | lateraloccipital | 0.163 | 0.00062 | 0.914 | 7.68E-19 | 0.0012 | 0.738 | 0.585 |
|  | parahippocampal | 0.153 | 0.00075 | 0.897 | 5.1E-28 | 0.0038 | 0.656 | 0.466 |
|  | rostralmiddlefrontal | 0.181 | 0.000028 | 0.881 | 1.74E-15 | 0.000053 | 0.706 | 0.568 |
|  | superiorfrontal | 0.128 | 0.0012 | 0.950 | 1.17E-14 | 0.0051 | 0.825 | 0.745 |
| Surface Area | Hemisphere | 0.166 | 0.00025 | 0.992 | 3.73E-24 | 0.0036 | 0.963 | 0.925 |
|  | entorhinal | 0.243 | 0.0000047 | 0.737 | 6.15E-09 | 0.0039 | 0.437 | 0.289 |
|  | inferiorparietal | 0.189 | 0.000085 | 0.761 | 2.43E-11 | 0.000055 | 0.504 | 0.325 |
|  | isthmuscingulate | 0.167 | 0.00028 | 0.729 | 3.5E-10 | 0.00015 | 0.397 | 0.176 |
|  | superiortemporal | 0.193 | 0.000014 | 0.844 | 2.64E-18 | 0.000069 | 0.591 | 0.361 |

## Discussion

In the largest ever analysis of asymmetry of cerebral cortical structure, we applied a meta-analytic approach to brain MRI data from 17,141 healthy individuals from datasets across the world. The findings revealed substantial inter-hemispheric differences in both regional cortical thickness and surface area, and linked some of these asymmetries to sex, age, and ICV. Handedness was not significantly associated with cortical asymmetries. While previous findings are based low hundreds of participants and different methodological approaches, this study of more than 17,000 participants is a major step forward in achieving a more accurate description of the typical asymmetries of the human brain, as well as variation in these asymmetries and some key individual differences factors which affect them. Moreover, with two independent pedigree datasets (i.e., GOBS and HCP), we revealed that several regions showed significant heritability of asymmetry measures.

### Cortical thickness

Regions with significant leftward asymmetry in thickness (i.e., left > right) were identified mainly in the frontal cortex, as well as the primary sensory, superior parietal, and medial temporal cortices, while rightward asymmetry was prominent in the posterior cortex, including lateral and medial parts of the temporal, parietal and occipital cortices. This striking asymmetry pattern along the fronto-occipital axis (see Figure 4) is similar to that reported by Plessen et al. (2014) and may be related to the “Yakovlevian torque”, i.e. the frontal/occipital bending in the human brain (34). Specifically, the torque refers to the phenomenon of crossing of the interhemispheric fissure by one hemisphere into the domain of the other. The frontal and occipital bending are the main twisting effects of the torque in opposite directions, with right frontal bending to the left, and left occipital bending to the right (60). At the population level, we found that the frontal regions showed leftward asymmetry in cortical thickness, while the occipital regions showed rightward asymmetry.

There were some inconsistencies when comparing our results with previous studies. For example, in 215 healthy participants, Plessen et al. (2014) observed a leftward asymmetry in the inferior frontal cortex, which includes Broca’s area in the inferior frontal gyrus. The authors suggested that this might correspond anatomically with the functional asymmetry for expressive language in these regions, as has been reported on the basis of brain lesion studies and functional neuroimaging studies (61–63). However, this interpretation should be considered with caution in light of a recent study on cortical thickness asymmetries with 250 adults showing an opposite direction of asymmetry (rightward) in this region (44). In the present study, with a much larger sample size, we failed to detect any cortical thickness asymmetry in this region (i.e., the pars opercularis and pars triangularis of the inferior frontal gyrus, uncorrected *p* >0.45). Another difference with previous findings concerns the supramarginal gyrus, which showed a strong leftward asymmetry in Plessen et al. (2014), but no asymmetry in two other studies (43, 44), and also not in the present study. This indicates an absence of population-level lateralization in cortical thickness in the supramarginal gyrus, and again underlines the value of the present study in achieving a more accurate characterization of the average anatomical brain laterality.

There are several issues that may contribute to discrepancies of our present results with these previous studies, including the large sample size that we used, as well as the worldwide population. Varying demographic factors, such as sex and age, across the various previous studies might also have played an important role. In the current study, we identified several regions showing significant effects of these factors on the asymmetry of cortical thickness. For sex, notable effects were observed in the medial temporal regions, including the parahippocampal gyrus (more leftward in males) and the entorhinal cortex (more rightward in females), while mixed results have been obtained in previous studies (46, 50). Considering the critical roles of these two regions in visuospatial processing and spatial navigation (e.g., (64, 65)), these sex differences may be related to the tendency for slight male advantage on spatial tasks (66–68). In contrast to Plessen et al. (2014), we found no sex differences in cortical thickness asymmetry of core regions of the language network, including the pars opercularis and pars triangularis of the inferior frontal gyrus (the Broca’s area), the transverse temporal gyrus (the Heschl's gyrus), and the supramarginal gyrus (uncorrected *p* > 0.05). These results are consistent with two other studies (43, 50), and indicate that subtle sex differences in the performance on language tasks and language lateralization (67) cannot be linked to sex differences in cortical thickness asymmetry of these regions.

In terms of age effects, when limiting our analysis to only the datasets with an age range greater than 20 years, we found a significant correlation between age and the overall hemispheric asymmetry in cortical thickness (i.e., increasing age correlated with more pronounced leftward asymmetry), which was mainly contributed by the superior temporal gyrus. This finding is consistent with previous studies (46, 47), though we did not detect age effects in other regions reported by Plessen et al. (2014). Brain size is another factor that can affect functional organization (69, 70). In the present study, we found a significant effect of ICV on the overall asymmetry in cortical thickness, such that the leftward asymmetry in cortical thickness increases in larger brains. This effect was the most pronounced in the inferior parietal gyrus and the insula. Our findings on ICV are in accord with the hypothesis that asymmetries increase in larger brains, which might relate to the increased inter-hemispheric distance and transfer time in larger brains (71).

No detectable association of handedness with cortical thickness asymmetries was found, even in this unprecedented sample size. This negative result is consistent with one recent study (45), while in other studies handedness has not been included as a variable of interest (43, 46, 47, 50). It remains possible that handedness is associated with asymmetry measures of other structural metrics and/or in more narrowly defined regions. However, it is clear from the present results that left-handedness does not involve any broad or substantial alterations of cortical thickness asymmetry.

### Surface area

Regarding surface area, population-level asymmetry was generally more prominent compared to that of cortical thickness. A large majority of regions (91.1%) showed significant asymmetry in surface area, although with no obvious directional pattern (leftward or rightward) affecting multiple neighboring regions, or along the anterior-posterior axis, as we observed for thicknesses. The present study detected some similar asymmetry patterns of surface area to those of two previous studies (44, 49). Specifically, consistent results included leftward asymmetry of the superior frontal gyrus, the postcentral gyrus, supramarginal gyrus, and the entorhinal cortex, and rightward asymmetry in the caudal anterior cingulate cortex and the middle temporal gyrus (44, 45, 49, 50). The leftward asymmetry of surface area in the supramarginal gyrus is consistent with the widely-observed volume asymmetry in the Perisylvian regions, which is related to an asymmetrical shift caused by the brain torque (33, 37, 38, 45, 72).

We identified several additional regions that are asymmetric in terms of surface area, not previously described. Among these regions, two language-related regions, including the opercular part of the inferior frontal gyrus (the posterior part of Broca's area) and the transverse temporal gyrus ( Heschl's gyrus) showed the largest leftward asymmetries. Based on these findings, the asymmetry of surface area (rather than cortical thickness as suggested in Plessen et al., 2014) may correspond anatomically with language lateralization in these regions, although further study is needed investigating both structure and function. Moreover, we found two other language-related regions showing strong asymmetry in the opposite direction (rightward), including the triangular part of the inferior frontal gyrus (the anterior part of Broca’s area) and the inferior parietal gyrus. Taking these observations together, it appears that the structural basis of functional language lateralization is more complex than previously thought. For example, as mentioned above, for Broca’s area, one of the most well-established areas for language function and language lateralization, while we did not detect asymmetry in terms of cortical thickness, we indeed observed strong asymmetry in surface area within this region. Moreover, the asymmetry was in different directions in two sub-regions of this area: leftward for the posterior part and rightward for the anterior part. These findings may be closely related to distinct roles of these two sub-areas in language functions: these two sub-regions are involved in, respectively, phonology and syntax, related to their distinct connections with areas in inferior parietal and temporal cortex (73, 74). Thus, these findings suggested that the opposite directions of structural asymmetry affecting regions within one network or within one functional area might reflect different functional involvements of each component region. Future studies with both structural and functional data in same participants may help link the structural asymmetries to functional asymmetries in the human brain.

The effects of biological factors on surface area asymmetries were more prominent than on thickness asymmetries. Very few previous studies have reported sex effects. Kang et al. (2015) found no sex differences in asymmetries for surface areas in 138 young adults, while Koelkebeck et al. (2014) only reported a male > female effect for the asymmetry of surface area at the overall hemispheric level in 101 healthy individuals. We also found that males, on average, showed more rightward asymmetry in overall surface area, compared with females, which is consistent with Koelkebeck et al. (2014). We additionally observed a number of regionally specific effects, among which surface area asymmetry in the superior frontal gyrus showed the strongest relation to sex (i.e., males showed more leftward asymmetry in surface area in this region compared to females).

In terms of age, when including only those datasets with an age range of more than 20 years, we found a weak positive correlation between age and the asymmetry of surface area of the entorhinal cortex, that is, the leftward asymmetry of this region was slightly greater in older participants. As far as we are aware, no previous studies have reported possible age effects on the asymmetries of surface area, except one that showed no significant results in 101 participants (44). Note that, in our analyses for either sex or age effects, ICV was included as a covariate to obtain sex- or age-specific effects. In terms of ICV effects themselves (correcting for sex and age), no significant effect was found on the overall asymmetry of surface area, but a number of regionally specific effects were revealed. Specifically, positive effects (increased leftward/decreased rightward asymmetries with ICV) were observed mainly in medial regions such as the anterior cingulate gyri, while negative effects (decreased leftward or increased rightward asymmetries with ICV) were seen in spatially diverse locations, including the posterior cingulate gyrus, the insula, and the caudal middle frontal gyrus. It has been suggested that increased brain size might lead to the development of additional sulci (50), which could impact on regional asymmetries as assessed with the FreeSurfer atlas-based approach.

No significant associations of handedness with cortical surface area asymmetries were found. This negative result is consistent with two recent studies (30, 45), while in other studies handedness has not been included as a variable of interest (44, 49, 50). Note that our results did include several nominally significant handedness associations in several temporal regions, including the fusiform, parahippocampal, and superior temporal gyri. However, these effects did not survive correction of multiple comparisons, and further studies are needed to confirm or refute these associations.

### General discussion

Our findings bear on the relationship between asymmetry of cortical thickness and surface area. Previous studies have suggested that thickness and surface area are evolutionarily, genetically, and developmentally distinct (40, 75), and that therefore separate consideration of these aspects of cortical anatomy is important (e.g., (76)). With a large MRI twin sample, Panizzon et al. (2009) showed that, although average cortical thickness and total surface area are both highly heritable (>0.80), they are essentially unrelated genetically (genetic correlation = 0.08). This genetic independence of cortical thickness and surface area was also found in a large extended family study (76). These results suggest relative independence of the two surface-based measures, and potentially therefore their asymmetry patterns. Data from two recent studies has indeed indicated that the asymmetry measures of cortical thickness and surface area are relatively independent at the overall hemispheric level (44, 45). With our larger sample size in the present study (including the BIL&GIN dataset used in Maingault et al., (2016), we confirmed a lack of correlation across regions between the asymmetries of thickness and surface areas, which further supports their independent natures. Moreover, by including data on participants’ sex, age, handedness, and ICV, our findings further elaborated the largely independent nature of regional area versus thickness variability. That is, no overall correlations were found between the effects of these factors across regions, on cortical thickness and surface area. Note that, when zooming in on some individual regions, there may be identifiable relations between thickness and surface area asymmetries, such as reported for the fusiform gyrus and the cingulate cortex (44, 45, 49), although further investigation is needed. In future studies of cortical asymmetry, the simultaneous investigation of both cortical thickness and surface area will be important. For example, this may be necessary in order to approach the genetics of brain asymmetry (30) and its links with functional lateralization (e.g., language lateralization) (77).

With the pedigree datasets from the GOBS and HCP, we revealed that several regions showed significant heritability of their asymmetry measures, while the general distributions of heritability estimates across the cortical regions were quite different for thickness and surface asymmetries (Supplemental Information S8). These data on heritability will be useful in targeting future studies of brain laterality with, for example, genome-wide association scanning aimed at identifying genes involved. Interestingly, cortical asymmetry of the human brain may also be associated with inter-hemispheric differences in gene expression (78, 79). Moreover, besides the directional asymmetry (DA) studied in this study, fluctuating asymmetry (FA), defined as the distance from the population-level mean directional asymmetry, reflects environmental influences during development but shows significant heritability (80). In future ENIGMA genome-wide association studies of brain asymmetries, both indexes may be included as the phenotypes.

Our data revealed extensive variability in cortical asymmetry across participants and samples. Besides sex, age, brain size, handedness, and heritable effects, further studies on individual variability are needed, from the perspective of cognitive and neuropsychiatric disorders. Some disorders, such as dyslexia (11), Alzheimer’s disease (12), ADHD (13), psychotic disorders (14–17), autism (18), and mood disorders (19, 20), may be associated with abnormal cortical asymmetries, though these complex links have not been fully explored. Asymmetry measures may even be more accurate than unilateral cortical measures to distinguish healthy controls from patients in some contexts (81), suggesting the potential for cortical asymmetry to be used as an important biomarker. In this respect, the findings in this paper provide a reference for cortical asymmetry in healthy populations, which may help for further understanding the nature of these disorders in future studies. In fact, studies of cortical asymmetry in several disorders are currently underway within the ENIGMA consortium, using the same methodology as used here.

Regarding handedness, it is interesting to note that paleoneurologists have attempted to use skull endocasts to assess cerebral asymmetries and to infer the evolution of handedness in hominins (82). Since we found no significant association between brain anatomical asymmetries and handedness, our analysis does not support the use of indirect measures of brain anatomy to infer the handedness of individuals.

### Limitations and future directions

This study has several limitations which could be overcome in future studies. First, for age effects, the cross-sectional study design limits the interpretation of results. Longitudinal studies should ideally be performed to support the findings and provide the normal course of development and aging of brain asymmetry.

Second, when combining already collected data across worldwide samples, data collection protocols are not prospectively harmonized. Imaging acquisition protocols and handedness assessments therefore differed across studies, which resulted in possible sources of heterogeneity. On the other hand, this heterogeneity can be taken as an advantage of our approach, in the sense that our findings are representative of the real-world diversity of MRI acquisition currently in use in the field, and not limited to a single lab-specific protocol.

Third, we note that variability of asymmetry in surface area across samples was relatively lower than that of asymmetry in cortical thickness, at both the global hemispheric and regional levels (see Supplemental Information S5). The relatively consistent asymmetry in surface area across datasets might be, to an extent, driven by the same parcellation scheme having been used across all samples. The potential impact of this issue is an important direction for future studies. In addition, the region-based approach is necessarily limited in terms of spatial resolution, related to the number of cortical regions defined. A vertex-wise approach combined with cross-hemispheric registration methods could be used in future cortical asymmetry studies (45, 50, 83).

Finally, besides the directions of the asymmetries, the present study provided the exact effect size distributions for each region with a very large sample size. The results can act as a guide and provide a reference normative resource for future studies of cortical asymmetry. For example, with the population-level effect sizes, researchers can estimate sample sizes required to detect specific effects of interest. Researchers can query the meta-analysis summary statistics with the query tool (http://conxz.github.io/neurohemi/).

### Summary

In summary, we showed that diverse regions of the human cerebral cortex are asymmetrical in their structural features (i.e., cortical thickness and surface area) with different effect sizes, and that the asymmetry patterns are different between cortical thickness and surface area. Moreover, we showed widespread effects of several biological factors (e.g., sex, age and ICV) on the cortical asymmetries, but found no significant handedness effects. Finally, we revealed that the human brain is composed of regions with significant heritability of the asymmetry characteristics. This study not only contributes to the understanding of human brain asymmetry in the healthy population, but also provides informative data for future studies of the genetics of brain asymmetry, and potentially abnormal brain asymmetry in cognitive and neuropsychiatric disorders.

## Methods

### Datasets

The primary datasets used in this study for large-scale meta-analysis were from members of the Lateralization Working Group within the ENIGMA Consortium (57). There were 99 independent samples, including 17,141 healthy participants from diverse ethnic backgrounds. Samples were drawn from the general population or were healthy controls from clinical studies. Figure 1 summarizes the age ranges and sample sizes of each dataset (for more details on each sample, see Supplemental Information S1). All local institutional review boards permitted the use of extracted measures of the completely anonymized data.

**Figure. 1.**
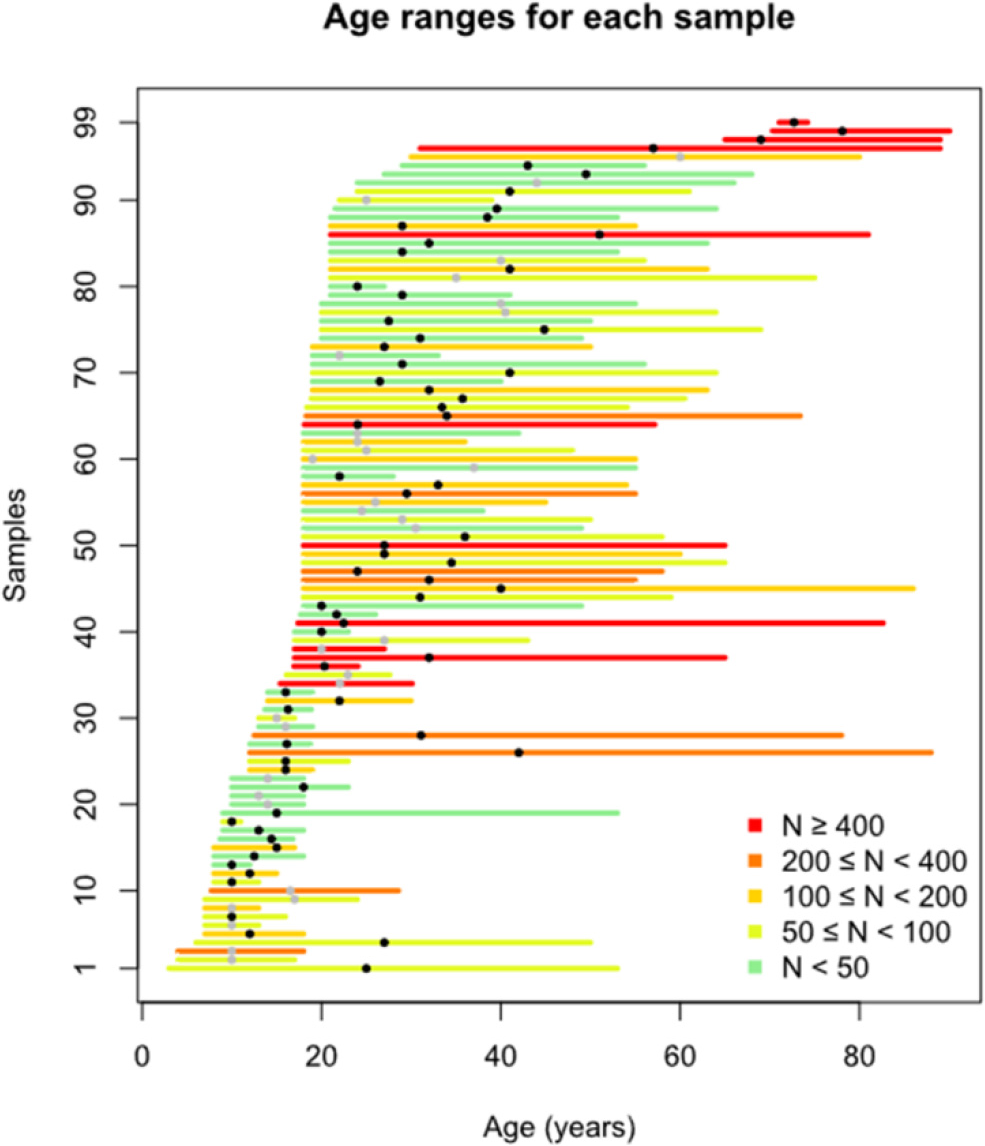
The age ranges and sizes of each sample. Each line covers the age range of an individual sample, with different colors indicating the sample sizes (see color key in figure). The position of the gray/black dot on each line indicates the median age of that sample. Black dots indicate samples with handedness information available. For more details, see Supplemental Information S1.

Handedness was known for a subset of the participants. The method of assessment varied across samples (see Supplemental Information S2). An ambidextrous category was not included, resulting in 827 left-handed and 11,237 right-handed participants in total.

Two additional datasets were used to estimate heritability of asymmetry measures, i.e. the GOBS dataset and the HCP dataset. GOBS is a family study comprising 1,443 individuals with MRI data (836 females), aged between 18 and 85 years at the time of scanning (59). All GOBS subjects are Mexican Americans and belong to pedigrees of varying sizes (the largest pedigree has 143 members). The HCP is a large-scale project comprising 1,113 individuals with MRI data (606 females, age range 22-37 years at the time of scanning) of varying ethnicities (http://humanconnectome.org/). The HCP contains 143 monozygotic twin pairs and 85 dizygotic twin pairs, as well as unrelated individuals.

### Image Acquisition and Processing

Structural T1-weighted MRI scans were acquired and analyzed locally. Images were acquired at different field strengths (1.5 T and 3 T). The images were analyzed using the fully automated and validated segmentation software FreeSurfer (version 5.3 for 91 of 99 samples). Segmentations of 68 (34 left and 34 right) cortical regions based on the Desikan-Killiany atlas (84) were statistically evaluated and in some cases visually inspected for outliers. Regional measures of cortical thickness and surface area were extracted for each participant. In addition, two hemisphere-level measures (average cortical thickness and total surface area), as well as the ICV were obtained. Quality control and data analysis were performed following standardized ENIGMA protocols (see http://enigma.ini.usc.edu/protocols/imaging-protocols/). In addition, following Guadalupe et al. (2016), we performed several checks to assess potential errors in the left-right orientation of the data. Further details for the datasets and orientation checking can be found in Supplemental Information S2.

### Within-dataset Analyses

For each dataset, descriptive and statistical analyses of the asymmetries in both cortical thickness and surface area were performed at each participating site using a single script in the R language, based on unified table-formatted data. For each global hemispheric or regional measure, an asymmetry index (AI) was defined as (L-R)/((L+R)/2), where L and R are the corresponding thickness or area measures on the left and right hemisphere, respectively. Thus, positive and negative AI values indicate leftward and rightward asymmetry, respectively. To exclude possible outliers in measures of cortical thickness, surface area, or AIs, we followed Guadalupe et al. (2016) and used an adaptive threshold (SDthre) depending on each dataset’s sample size: N < 150, SD_thr_ = 2.5; 150 ≤ N ≤ 1000, SD_thr_ = 3; N > 1000, SD_thr_ = 3.5.

Statistical tests were run for each hemisphere-level or regional measure separately. Paired t-tests were used to assess inter-hemispheric differences. Cohen’s d was calculated based on each paired t-test result to estimate the effect size of population-level asymmetry. All differences between sexes (−1=females, 1=males) were assessed with linear regression models adjusted for age, age^2^, and brain size (as indexed by ICV). Cohen’s d was calculated to estimate the effect size for each comparison. Furthermore, we examined the age effects on the AIs in cortical thickness and surface area, adjusting for sex and ICV. Similarly, we examined associations between ICV and the AIs in cortical thickness and surface area, adjusting for age, age^2^, and sex. In addition, if handedness information was available, AI differences between handedness groups (−1=left; 1=right) were assessed with linear regression models adjusted for all the other covariates. For each analysis above, additional covariates of scanners were included when more than one scanner was used at one site.

### Meta-analyses

All regression models and effect size estimates were fitted at each participating site separately. We then combined the output statistics from each dataset using inverse variance-weighted random-effect meta-analyses (85) with the R package *metafor*, version 1.9-9. This method tests one overall effect, while weighting each dataset’s contribution by the inverse of its corresponding sampling variance. Thus, unlike fixed-effect meta-analysis, this method takes into account variability across different studies.

A Cohen’s d effect size estimate of population-level asymmetry (hemispheric difference) was obtained using a random-effect meta-analysis model for each region and each cortical measure (cortical thickness and surface area). Note that including results based on too few participants may reduce reliability, and therefore we only included datasets with a sample size larger than 15. In the meta-analysis, heterogeneity of each effect was assessed via the I^2^ value, which describes the percentage of total variation across studies that is due to heterogeneity rather than chance. I^2^ values of 25%, 50%, and 75% indicate a low, moderate, and high heterogeneity, respectively.

Similarly, Cohen’s d estimates of sex and handedness effects on asymmetry were obtained by meta-analysis for each cortical region. Again we only included samples with at least 15 participants per group. For meta-analyses of the correlations of asymmetries with age or ICV, predictors were treated as continuous variables, so that effect sizes were expressed as partial-correlation *r*. For metaanalyses of age effects, we only included samples with a minimum 5-year range in the initial analysis, and in a subsequent analysis we further restricted to samples with age ranges larger than 20 years, to better capture the age effects in our data.

In this study, we report uncorrected *p* values with a significance threshold determined by Bonferroni correction for multiple comparisons within each separate meta-analysis (i.e. correction separately for analysis of 34 regional surface area asymmetries and 34 regional thickness asymmetries, *p* = 0.05/34 = 0.00147). No correction was done for global hemispheric measures of asymmetry.

### Moderator analyses with meta-regression

Meta-regressions were performed to evaluate the potential moderating effects on meta-analysis effect sizes. We tested whether moderating factors, including median age, median ICV, sex ratio, handedness ratio, and MRI scanner field strength (3T, *N* = 63 datasets versus 1.5T, *N* = 28 datasets), influenced the effect size estimates across datasets in the meta-analyses. Each moderator variable was separately included as a fixed effect predictor in the meta-regression model. All statistical analyses were conducted using the R software package *metafor*, and Bonferroni correction was applied for multiple comparisons (*p* < 0.00147; see above).

### Heritability estimation

Furthermore, we estimated the heritability of asymmetries in cortical thickness and surface area, firstly using the GOBS dataset (*N* = 1,443; see ‘Datasets’ above). For more details about this cohort, see McKay et al. (2014). Specifically, we estimated the narrow-sense heritability (i.e., the proportion of overall variance explained by additive genetic effects) of each AI using variance-components analysis (86). Briefly, each AI was entered as a dependent variable into a linear mixed-effects model, which included fixed effects of age, sex, and ICV, and a random effect of genetic similarity, whose covariance structure was determined by the pedigrees. We refer to these as univariate polygenic models. Second, we estimated the genetic correlations between left and right thickness/area measures by extending the univariate polygenic models to incorporate two traits at once; these bivariate polygenic models simultaneously estimate the heritability of left and right paired measures, along with their genetic correlation (which indicates the extent to which their variation is influenced by the same genetic factors).

In principle, family-based analysis such as that using GOBS can conflate shared environmental effects with heritable effects, whereas a twin design is robust to this. Therefore, we also performed heritability analysis in the HCP cohort (N = 1,113; see ‘Datasets’ above). HCP is a large-scale project which includes monozygotic and dizygotic twin pairs, as well as unrelated individuals. Precisely the same analyses were conducted in this second cohort.

## Collaborators

Members of the Karolinska Schizophrenia Project (KaSP):

## Acknowledgements

Funding information for each site is available in a separate Acknowledgements document. All sites within the ENIGMA-Lateralization working group are very grateful to the participants for their generosity of time and willingness to participate in each of the collaborating studies.

## Compliance with ethical standards

All participating studies were approved by the local ethical committee. Informed consent was obtained from all participants involved. The authors declare no conflicts of interest except for the authors listed in Conflicts of interest.

